# Novel software for automated morphometric analysis of stented arteries

**DOI:** 10.1101/2020.01.30.927459

**Authors:** Aviel Resnick, Bahman Hooshdaran, Benjamin B. Pressly, David T. Guerrero, Ivan S. Alferiev, Michael Chorny, Robert J. Levy, Ilia Fishbein

## Abstract

Morphometric analysis of explanted arteries remains the gold standard in assessing pathological mechanisms of arterial disease, and the efficacy of therapeutic interventions. Software currently used for morphometric analysis (ImageJ, ImagePro, etc.) requires manual tracing of each region of interest by the user to obtain direct and derived data needed for analysis. Manual segmentation of borders between differently stained arterial compartments is time-consuming and prone to bias, even when the operator is blind to the treatment type. Here, we report newly designed morphometric software, floodSeg, which greatly aids analysis through semiautomated segmentation, followed by automated computation and data entry. The program includes tools for both rapid component selection, and precision user-controlled contour correction. In practice, segmentation occurs following the selection of seed points and corresponding threshold values, for each desired component. The flood fill algorithm is used to map out components, followed by Sklansky’s convex-hull algorithm to obtain the outer contours. If necessary, convexity defects can be overcome through manual point placement on top of existing points, and regeneration of the contour. floodSeg was tested using a set of non-uniformly stained stented rat arteries, and compared against manually obtained results. The accuracy of the resulting measurements was within the expected limit based on repeated manual measurements by the same operator, and did not exceed 3%. Most notably, the duration for data acquisition using floodSeg was less than 20% of the time required for manual measurements by an experienced operator. Thus, our contribution is an improvement on widely used software, with significant potential for application in a multitude of areas of pathology practice.

## Introduction

Microscopic evaluation of pathological processes is pivotal for diagnostics, treatment, and prognostication in clinical medicine, as well as an indispensable tool for medical research. A majority of contemporary microscopic studies include elements of morphometry, i.e., quantitative measurements of defined areas in a digital microscopic image. Several popular software packages, both commercial (ImagePro, MIPAR, Imaris, Lucia Image, SigmaScan) and freeware (Image J, NIH Image), provide the necessary graphical tools for tracing specific elements in digital images with subsequent computation of areas, perimeters, angles, distances, etc. A calibration option is typically available to convert the data from pixels to metric units. Depending on the specific application, these morphometric tasks are often extremely time consuming and prone to bias. Inter-and intra-evaluator variability is usually high in these studies (1, 2). Therefore, automated image analysis software is increasingly used to facilitate morphometric tasks and unify analysis criteria, especially between different laboratories working in oncology(3), nephrology(4), gastroenterology(5), orthopedic(6) and neurology(7) research. Furthermore, the USA Food and Drug Administration has recently determined substantial equivalence of certain automated image analysis implementations to manual analysis performed by trained pathologists (8), thus lifting regulatory barriers to the wide-spread use of automated image evaluation of clinical samples.

While the main clinical manifestations of atherosclerosis such as myocardial infarction and stroke remain the main causes of death worldwide, relatively minor efforts were made to develop specialized software for automated analysis of normal and diseased vasculature. The most probable explanation of this fact is that clinical samples of large and mid-sized arteries are very uncommon, and are not expected to have any direct impact on clinical practice. However, the research focusing on animal models of atherosclerosis and restenosis is burgeoning with more than 50,000 papers published during the past five years in PubMed-referenced journals. Although many modalities such as angiography, ischemic and inflammatory biomarkers, are available to assess the atherosclerotic burden and extent of restenosis in diseased arteries, morphometric evaluation of arterial sections presents a gold-standard methodology providing a wealth of morphological and mechanistic information. Since the severity of disease varies across a given artery, multiple sections spaced 1 mm or less are typically examined and assessed in each animal, often resulting in several hundred histological sections per study.

Herein, we report on a novel computational solution for the semi-automated segmentation of specific anatomic compartments in stained sections of rat carotid arteries with in-stent restenosis. We show that the use of our new program, floodSeg, greatly facilitates morphometry by reducing the duration of analysis more than 5-fold, without compromising the accuracy of the measurements.

## Methods

### Animal model of arterial injury

Male Sprague-Dawley rats (400-450 gram) were anesthetized with ketamine and xylosine, and the common and external left carotid arteries were exposed(9). A 2-French Fogarty balloon catheter was introduced into the common carotid artery, via an arteriotomy made in the external branch, and the endothelial layer of the common carotid artery was denuded by three passages of the balloon inflated with 50 μl of saline. A 3-cm piece of a 1.1-mm OD Teflon tubing was introduced into the common carotid over the Fogarty catheter, and the catheter was exchanged for a stent-loaded angioplasty catheter. After deploying the stent in the mid-section of the common carotid artery at 12-14 atmospheres, the angioplasty catheter was withdrawn, the arteriotomy was tied-off both proximally and distally, the muscles and the skin were sutured, and the animals were recovered. The rats were euthanized 14 days after the surgery by CO2 asphyxia, and the stented segments of the common carotid artery were harvested and formalin-fixed. All animal procedures were carried out following the animal welfare regulations, and were pre-approved by the Institutional Animal Care and Use Committee.

### Tissue processing

Stent struts in formalin-fixed carotid segments were dissolved in a mixture of nitric and hydrofluoric acid, as reported by us before (10). Destented arteries were then routinely embedded in paraffin, sectioned (6 μm thick), and stained according to the Verhoeff-van Gieson method (9, 10). The stained sections were imaged at 40x magnification with a Nikon Eclipse 80i microscope equipped with Nikon DS-Ri1 CCD camera, and saved as digital images in JPG format with 1028×1240 resolution.

### Manual image analysis

Manual image analysis was conducted with the ImageJ (1.49v) program. The pixel-to-millimeter ratio was established, and used globally for the conversion of pixels into metric units.

Perimeters of the arterial lumen (LUp), internal elastic lamina (IELp) and external elastic lamina (EELp), as well as the areas confined within these lines (LUa, IELa, and EELa, respectively) were measured using a polygon selection tool. Additionally, areas originally occupied by the now dissolved stent struts and the minimal distances (MD) from these strut derived voids (SDV) to the lumen were measured (Fig. 1). These directly measured anatomical parameters were used to calculate indices of restenosis as follows:

1. *Lumen area* — *as measured (LUa)*;
2. *Neointimal area = IELa — LUa*;
3. *Percent of stenosis* 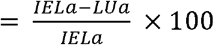;
4. *Distortion corrected percent of stenosis* 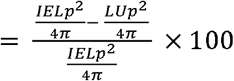;
5. *Neointima to media ratio* 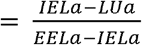;
6. *SDV corrected neointima to media ratio* 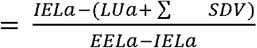, and
7. *Average neointimal thickness over the struts* 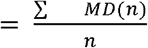.

**Fig. 1.**
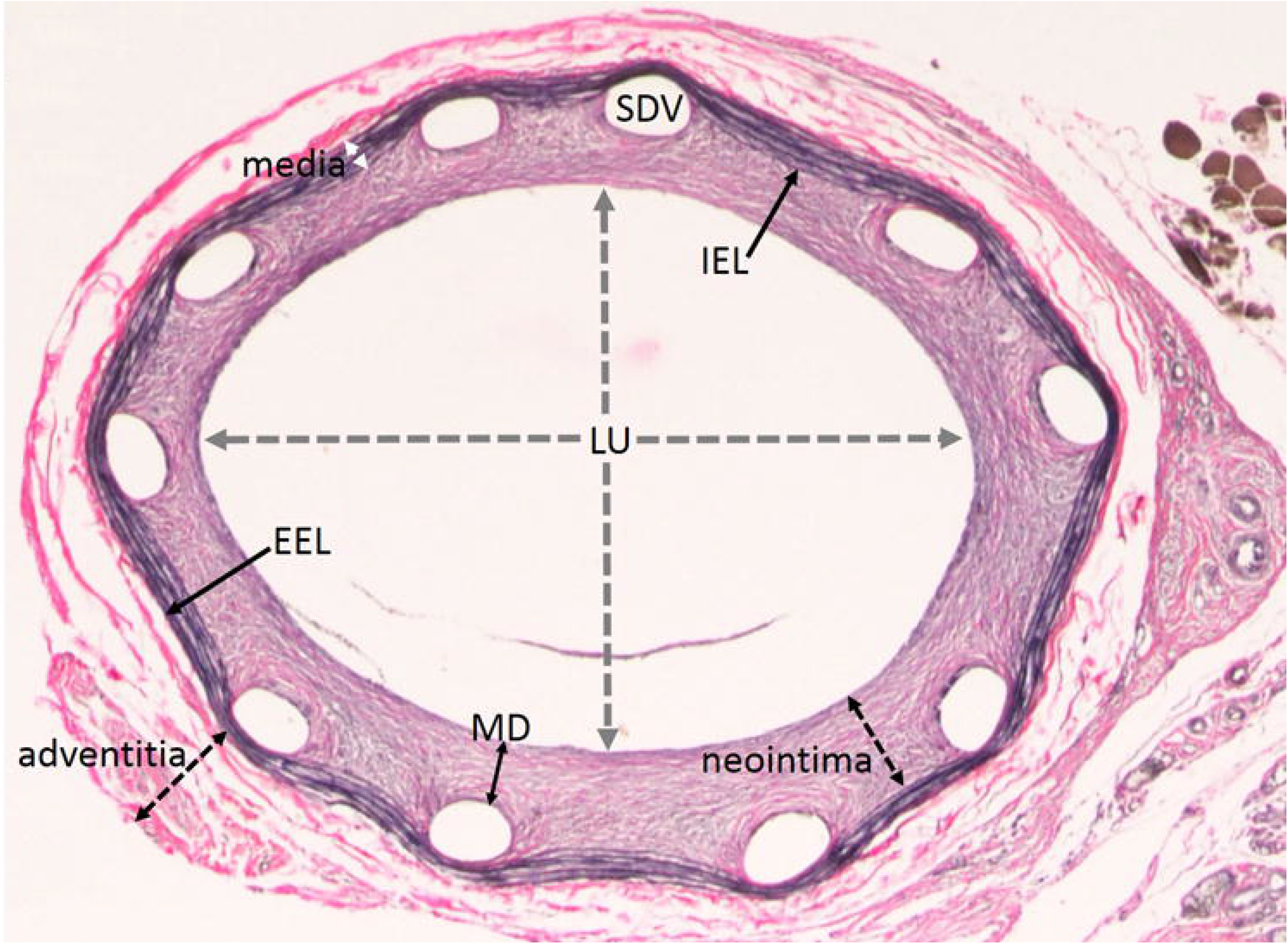
Annotated image of Verhoef-van Gieson stained section of a de-stented rat carotid artery (original magnification 40x). Arterial lumen (LU), neointima, media, adventitia, internal and external elastic laminas (IEL and EEL, respectively), strut derived voids (SDV) and minimal distance (MD) from SDV to LU are indicated with dashed lines and arrows.

### Computational methods

The current implementation of floodSeg is an open source python-based standalone application, with excellent potential as a platform for additional development. The philosophy behind floodSeg is rapid inside-out contouring, allowing for the calculation of additional values to be automated. Furthermore, this method allows for simple MS Excel and text based output. Since floodSeg was developed for morphometric analysis of vasculature, the “components” of interest are i) lumen; ii) neointima (Fig. 1), defined as the area located between the lumen and IEL; iii) media, defined as the area located between IEL and EEL, and iv) the struts-derived voids. It is important to note that given the inside-out method, only the outer edge contouring is of interest (e.g the EEL for the media component, or IEL for the neointima.) Programmatically, the components are stored in a dictionary, with string keys for the component names, and tuple values containing the area and arc length for each key. For each component starting with the innermost, an appropriate seed point should be selected by the user; a color average of the stained section often works well. The seed point is passed to a dedicated class which utilizes the OpenCV implementation of the flood fill algorithm to calculate the connected component within a color and brightness threshold of the seed. The outer contour of the component is then identified, and displayed on top of the image in a contrasting color for user approval (Fig. 2B). A threshold slider above the image allows the user to make real-time adjustments, and the seed point can be continuously repicked, providing the user with a high degree of control. Once the contour is satisfactory, the comprising points are returned to the main class, however in practice the component consists of many interconnected contours (Fig. 3A), and often no single outer contour can be found. To overcome this, the convex hull of the points is found with Sklansky’s algorithm(11) (Fig. 3B). While imperfect, the solution proposed by Sklansky (11) is one of the most popular algorithms for this problem, and works correctly in the vast majority of cases. The inherent problem with finding the convex hull, however, is that concave components are incorrectly contoured. These “convexity defects” (Fig. 3B Inset) must be corrected, but since all points are carried over (Fig. 3C), the user only needs to modify the points of concern (usually < 5 additions or removals). The contour is then regenerated to include the corrected points (Fig. 3D Inset). From there, OpenCV’s contourArea and arcLength functions provide the desired values, which are passed back into the component dictionary. In this particular study, there were multiple derived values which were calculated pre-output, including percent stenosis and neointima-to-media (N/M) ratio. Additionally, the arc lengths of components were used to create “calculated” areas, which corrected for the imperfect circular shapes of the arteries caused by mechanical impacts during sample processing. By using the inside-out contouring method previously discussed, automation of calculation and exporting is far easier; coupled with the speed of flood-filling the components, our method significantly improves the efficiency of computer-aided morphometric analysis.

**Fig. 2.**
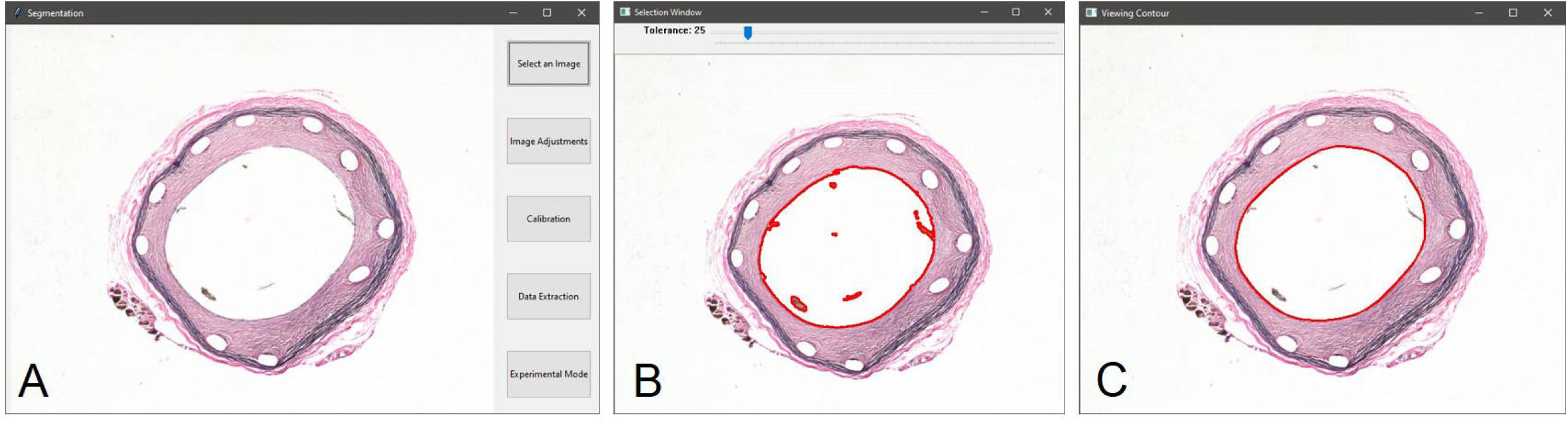
floodSeg applied to lumen segmentation. Representative screenshot images of A) destented Verhoeff-van Gieson stained rat carotid artery; B) same as A with the flood-fill algorithm applied to the lumen; C) same as B refined using the convex-hull algorithm.

**Fig. 3.**
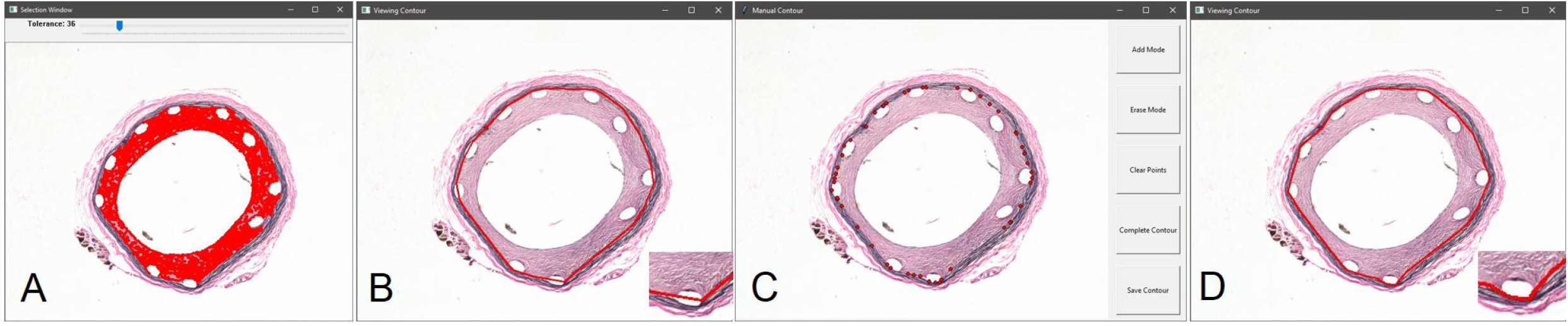
floodSeg applied to neointima segmentation. Representative screenshot images of A) flood-fill algorithm applied to the neointima of the same artery as in Fig. 2; B) same as in A refined using the convex-hull algorithm (convexity-defect attributed imperfections in the outer contour of the neointima are magnified in the inset); C) contour representation using latent constructing points from B; D) same as in B following user controlled correction of the outer contour (magnified in the inset).

### Comparison of manual and automated morphometry methods

Overall 7 images of Verhoeff-van Gieson stained de-stented rat carotid arteries originating from different animals were selected for analysis. The color hues, saturation, intensities and general brightness and contrast of the selected images were intentionally varied in order to validate the robustness of our system across the spectrum of staining results. Manual evaluation of each section was carried out 3 times, and the individual data for the directly measured and derived parameters were saved in MS Excel format. Additionally, the average time for each complete assessment of a single section was recorded. The floodSeg-based automated assessment of each section was carried out once by an operator blinded to the results of the manual study. Over the course of automated segmentation, several manual corrections were necessary to guide the process. The set of direct and calculated data obtained using floodSeg analysis was exported to MS Excel, and compared with 3 sets of manual data for potential outlier status (GraphPad; Grubbs’ ESD test)

## Results

Table 1 summarizes morphometric results obtained with ImageJ using manual tracing of the lumen, IEL, EEL, SDV and MD with subsequent calculation of restenosis indices, in comparison with floodSeg-acquired data. No floodSeg-based morphometry results were identified as outliers when added to the set of 3 manual measurements. In relative terms, the difference between the average of 3 manual measurements and floodSeg-generated values did not exceed 3%. Most notably, the total timing of floodSeg-assisted morphometry was 28 minutes, in comparison to the 154 min. manual procedure.

**Table 1.** Main morphometric results with manual (M) and floodSeg-automated (A) evaluation

| Artery | LU (mm<sup>2</sup>) |  | Neointima (mm<sup>2</sup>) |  | Neointima w/o SDV (mm<sup>2</sup>) |  | N/M ratio |  | SDV corrected N/M ratio |  | % stenosis |  | Distortion corrected % stenosis |  |
| --- | --- | --- | --- | --- | --- | --- | --- | --- | --- | --- | --- | --- | --- | --- |
|  | M | A | M | A | M | A | M | A | M | A | M | A | M | A |
| 1 | 0.590 | 0.591 | 0.266 | 0.264 | 0.224 | 0.227 | 2.742 | 2.722 | 2.309 | 2.340 | 31.075 | 30.877 | 35.291 | 35.749 |
|  | 0.591 |  | 0.268 |  | 0.226 |  | 2.913 |  | 2.457 |  | 31.199 |  | 35.851 |  |
|  | 0.593 |  | 0.259 |  | 0.216 |  | 2.643 |  | 2.204 |  | 30.399 |  | 35.147 |  |
| 2 | 0.394 | 0.395 | 0.385 | 0.383 | 0.346 | 0.349 | 3.702 | 3.648 | 3.327 | 3.324 | 49.422 | 49.229 | 50.416 | 51.458 |
|  | 0.397 |  | 0.381 |  | 0.342 |  | 3.699 |  | 3.320 |  | 48.972 |  | 51.743 |  |
|  | 0.396 |  | 0.384 |  | 0.345 |  | 3.728 |  | 3.350 |  | 49.231 |  | 51.847 |  |
| 3 | 0.444 | 0.442 | 0.357 | 0.363 | 0.319 | 0.329 | 2.746 | 2.814 | 2.454 | 2.550 | 44.569 | 45.093 | 46.986 | 46.950 |
|  | 0.444 |  | 0.362 |  | 0.324 |  | 2.850 |  | 2.551 |  | 44.913 |  | 46.429 |  |
|  | 0.445 |  | 0.359 |  | 0.322 |  | 2.740 |  | 2.458 |  | 44.652 |  | 47.656 |  |
| 4 | 0.468 | 0.466 | 0.359 | 0.360 | 0.324 | 0.329 | 2.943 | 2.927 | 2.656 | 2.675 | 43.410 | 43.584 | 40.296 | 39.967 |
|  | 0.463 |  | 0.362 |  | 0.326 |  | 2.919 |  | 2.629 |  | 43.879 |  | 40.059 |  |
|  | 0.467 |  | 0.358 |  | 0.322 |  | 2.934 |  | 2.639 |  | 43.394 |  | 40.136 |  |
| 5 | 0.393 | 0.391 | 0.335 | 0.342 | 0.278 | 0.282 | 3.564 | 3.638 | 2.957 | 3.000 | 46.016 | 46.658 | 47.530 | 49.045 |
|  | 0.391 |  | 0.342 |  | 0.285 |  | 3.800 |  | 3.167 |  | 46.658 |  | 49.665 |  |
|  | 0.392 |  | 0.337 |  | 0.280 |  | 3.510 |  | 2.917 |  | 46.228 |  | 49.483 |  |
| 6 | 0.463 | 0.461 | 0.376 | 0.375 | 0.333 | 0.333 | 3.357 | 3.409 | 2.973 | 3.027 | 44.815 | 44.856 | 49.249 | 46.060 |
|  | 0.460 |  | 0.373 |  | 0.330 |  | 3.161 |  | 2.797 |  | 44.778 |  | 45.122 |  |
|  | 0.467 |  | 0.373 |  | 0.330 |  | 3.657 |  | 3.235 |  | 44.405 |  | 43.671 |  |
| 7 | 0.384 | 0.391 | 0.341 | 0.339 | 0.287 | 0.279 | 3.279 | 3.645 | 2.760 | 3.000 | 47.034 | 46.438 | 48.288 | 48.652 |
|  | 0.389 |  | 0.345 |  | 0.289 |  | 3.876 |  | 3.247 |  | 47.003 |  | 43.943 |  |
|  | 0.396 |  | 0.332 |  | 0.277 |  | 3.952 |  | 3.298 |  | 45.604 |  | 54.030 |  |

## Discussion

The clinical definition of restenosis is a ≥ 50% angiographic narrowing of the stented segment when compared to the vessel diameter recorded immediately after stent deployment (12). Morphological criteria of restenosis are less explicitly defined, and are based on multiple morphometric indices, both directly acquired and calculated (13). In animal models, morphometric indices of restenosis reflect, in part, the experimental endpoints. Since the lumen diameter and percent of cross-sectional narrowing (percent of restenosis) are immediately linked to the blood flow across the stented segment of the artery and the downstream ischemic sequela, these indices are more suitable for clinically oriented translational research. The neointimal burden (neointimal area) and the thickness of neointima over the stent struts are of special interest for studies investigating the inhibiting effects of locally and systemically delivered therapeutics on the expansion of neointimal tissue, which is the main pathogenic component of in-stent restenosis.

The neointimal area, as a difference between the areas enclosed within the lines formed by the IEL and LU, is highly sensitive to the angle of sectioning which ideally would be perpendicular to the long axis of the artery. Since this is not always feasible, the neointima-to-media ratio, which is independent of the angle of sectioning, is often used instead of neointimal burden.

The volume occupied by struts or strut-derived voids are not always subtracted from the total neointimal area. This may be justified if the main focus of the study is on blood flow distal to the implanted stent, but should be avoided in experiments measuring neointimal response to therapeutic interventions.

The typical circular shape of vessels is often lost during the pre-embedding and paraffin embedding steps, especially in samples with relatively thin neointima. The extent of restenosis presented as neointimal area and neointima-to-media ratio is not affected by this mechanical deformation. However, the percent of restenosis changes as the arterial specimen deviates from its original circular form. In extreme cases, when the artery is completely flattened, thus obliterating the lumen, restenosis erroneously approximates a maximal 100%. To amend this problem, calculation of percent of restenosis based on conversion of perimeters of LU and IEL to areas of the respective circles is implemented using simple geometric considerations. This algorithm allows for the “restoration” of pre-deformation dimensions of the specimen, thus providing more reliable data which better reflect anatomical and physiological performance of the stented artery in vivo. In the current study, we determined multiple restenosis indices, including those discussed above, in order to establish the usefulness and robustness of floodSeg-based morphometry.

The floodSeg software was designed to be both modular and intuitive, acting as a toolbox for morphometric analysis of stent-implanted arteries. It provides multiple segmentation methods, including manual point placement, assisted flood-fill, and a fully automated neural network which is currently in development. Any method can be chosen depending on the shape and complexity of the section, and manual corrections can be made easily. The initial step is importing the image, followed by optional modifications (sharpness, contrast, and brightness), and inputting an optional pixel to metric value. The data extraction window provides a component overview, including statuses and options for adding/removing components. Once a component is selected, the user can choose to use either flood fill or manual point placement to obtain the outer contour. At any point, the “saved” contour can be viewed and modified manually. If a component is disconnected, for example multiple strut derived voids, the component can be split into multiple distinct parts all saved under one object. Once contouring is complete for all components, the user can export to MS Excel or a text file, which also triggers computation of derived values. In general, however, the list of desired components can be modified to fit any dataset. For example, if remodeling, rather than in-stent restenosis is the subject of study, the adventitial area (Fig. 1) can be determined by tracing the outer contour of the image.

Our method provides multiple distinct advantages over previously proposed solutions to computer-aided morphometric analysis. In the presence of image defects arising from foreign bodies, or other elements which interfere with sections of interest, the use of section hulls (solely outer contours) eliminates the need for user-correction. Alternative approaches, especially those based on pixel intensity(14), detect these defects as contrasting with the section they occupy, and exclude it from measurement without user correction. While our system similarly detects intra-sectional outliers, our interest in outer contours allows the system to ignore them, providing accurate results without correction. To the best of our knowledge, ours is the first application of convex-hulls to morphometric analysis. Furthermore, by maintaining a python dictionary of areas and arc lengths for each section, we are able to automate the computation of derived values such as percent stenosis and neointima-to-media ratio. Finally, the flexibility of our system lends itself to a multitude of applications, unlike hardcoded software dependent on specific color ranges.

We plan to further develop our method to include automatic computation of average neointimal thickness. Considering thickness as the average of shortest distances between SDVs and the lumen, the problem is reduced to finding the shortest path between known circular contours. Optimizations such as this are greatly aided by the floodSeg software, and we hope to further reduce the human intervention needed in accurate computer-aided morphometry.

In conclusion, based on our experience, a novel open-source software for image analysis, floodSeg, effectively facilitates morphometry of stented arterial segments, and significantly reduces the time and effort required.

